# Cleaner gobies can solve a biological market task when the correct cue is larger

**DOI:** 10.1101/2024.04.17.589842

**Authors:** Maddalena Ranucci, Mélanie Court, Beatriz P. Pereira, Daniele Romeo, José Ricardo Paula

## Abstract

Animal cognition is deeply influenced by interactions with the environment. A notable example of sophisticated cognition in the animal kingdom is described by the mutualistic relationship between cleaner fish and clients, where decision-making processes play a pivotal role in partner choice and fish survival. In this context, while extensive research has explored the cognition of the cleaner wrasse *Labroides dimidiatus*, its Caribbean counterparts, *Elacatinus spp*., have been comparatively inadequately evaluated. In this study, we used plexiglass plates as surrogates for clients and assessed the ability of cleaner gobies, *Elacatinus oceanops*, to solve a biological market task where prioritising an ephemeral food plate over a permanent one would double the food reward. We varied cue-based decision-making using both ecologically relevant cues (plate size and colour) and non-relevant ones (presentation side). Additionally, we tested their capacity for reversal learning, an indicator of complex cognitive abilities. Notably, cleaner gobies were able to solve the biological markets task when the distinguishing cue was a larger plate size. Given that these gobies tend to prioritise larger predatory clients in nature, our results align with their natural inclination. Furthermore, considering these gobies were bred in captivity and never experienced cleaning interactions in their lifetime, our data might suggest that their intrinsic cognitive abilities are shaped by evolutionary pressures rooted in their ecological roles. In essence, even in the absence of direct ecological interactions, innate cognitive abilities in cleaner gobies seem to be deeply influenced by evolutionary forces tied to their natural ecological functions. However, their inability to solve the same task involving other cues, may be influenced by factors such as captivity-reared fish or limitations in the experimental design. Further research, including wild individuals, is essential to elucidate the cognitive abilities of the studied species and its implications in the ecological context and evolutionary history.

## 1 Introduction

The survival and reproductive success of animals within their ecological niche are shaped by intricate interactions among behaviour, ecological pressures, and natural selection (Duckworth, 2009). Evolutionary changes in individual fitness are primarily driven by behavioural mechanisms (Bateson and Gluckman, 2012). Simultaneously, an individual’s behaviour is moulded by both biotic and abiotic environmental factors (Healy and Braithwaite, 2000). In this context, cognitive abilities, defined as the capacity to process, internalise and act on information acquired from the environment (Shettleworth, 2001), stand as a crucial factor for animal survival and reproduction success (Sol *et al*., 2007; Cole *et al*., 2012; Ashton *et al*., 2018). Cognitive performance, which relies on brain functions with energetic demands, is proposed to vary due to phenotypic plasticity in response to changes in energy availability and environmental demands (Maille and Schradin, 2017). Such variation in cognitive performance may impact decision-making processes, including partner choice and resource allocation. For instance, spatial discrimination abilities are more developed in environments where food availability is not immediate (White and Brown, 2015; Carbia and Brown, 2019). Conversely, in environments where predator threat is constant, cognitive skills related to perception and rapid response became more pronounced (Brown and Braithwaite, 2005). The complex dynamics of social groups within animal communities also require sophisticated cognitive skills to recognise individuals, understand social hierarchies and interpret communicative signals (Fernald, 2017; Kappeler, 2019).

Understanding animal cognition requires investigating innate cognitive abilities and adaptations influenced by the ecological context and evolutionary processes (Cauchoix and Chaine, 2016; Szabo *et al*., 2022). Animals’ innate abilities can vary from basic perceptual processes to complex problem-solving abilities (Rowell, Pillay and Rymer, 2021). Investigating this aspect offers the basis for comprehending how animals interact with their environment and prompts questions about the adaptability and plasticity of cognitive abilities.

In this regard, coral reefs present distinctive challenges, requiring animals to develop adaptative strategies and specialised skills for survival (Dixon *et al*., 2015; Picq, McMillan and Puebla, 2016; van Oppen *et al*., 2018). While often underestimated, fishes display remarkable cognitive capabilities essential for their survival and success, particularly in the context of avoiding predators and increasing survival (Brown and Braithwaite, 2005). These cognitive skills became particularly evident in mutualistic relationships, such as the one established between cleaner fish and their clients, where the ability to prioritise specific clients over others suggests cognitive sophistication (Triki *et al*., 2019).

Cleaning symbiosis is one of the most notable mutualisms in nature, and it plays a crucial role in maintaining ecosystem dynamics on coral reefs (Côté, 2000). Reef fishes are usually infested by parasites that can cause irritation or pose a risk of diseases. To eliminate these parasites, reef fishes referred to as ‘clients’ approach the ‘cleaning station’, a relatively fixed territory in the coral reef, allowing cleaner fish to remove parasites and dead skin from their bodies (Grutter, 1999; Vaughan *et al*., 2017). Through this interaction, there is an exchange of commodities between species where the cleaner fish gains a food source, and the client gets rid of parasites (Bshary, 2003).

In the Caribbean, cleaner gobies *Elacatinus spp*. provide an essential function to the ecosystem through cleaning mutualism (Côté and Soares, 2008; Vaughan *et al*., 2017). Here, cleaner gobies engage in with a wide variety of clients, including potentially threatening predators (Soares, Cardoso and Côté, 2007). Despite evidence of partner choice in natural observation (Côté and Soares, 2008; Soares *et al*., 2008, 2013; Dunkley *et al*., 2019), the cognitive abilities of cleaning gobies have been poorly investigated.

Partner-choice-derived cognition has been widely tested in cleaner fishes using a learning paradigm based on a fish’s capacity to solve discriminatory two-choice tasks, referred to in this context as partner-choice task, (Noë and Hammerstein, 1994; Mazzei et al., 2019; Triki et al., 2019; Wismer et al., 2019; Truskanov et al., 2021; Bshary and Noë, 2023), inspired by the principles of biological market theory (Noë and Hammerstein, 1995; Hammerstein and Noë, 2016; Noë, 2016). This theory provides insight into how organisms make decisions based on trading and cooperation observed in human economic markets. Within the context of cleaning mutualism, where the clients act as consumers and the cleaners as service providers, these ecological interactions resemble transactions in a marketplace(Hammerstein and Noë, 2016). These services are analogous to goods or resources of a traditional market, with clients benefiting from removing ectoparasites. Cleaner gobies prioritise their clients based on specific cues determining the order in which cleaning services are provided(Noë and Hammerstein, 1994; Noë, 2016; Triki et al., 2019).

In our experiment, we applied this principle to assess the cognitive abilities of cleaning gobies. The task required them to make decisions mirroring their ecological context. (Salwiczek *et al*., 2012; Gingins and Bshary, 2016; Triki *et al*., 2019; Wismer *et al*., 2019) Specifically, the fish were presented with two plates and were asked to choose between one of them, giving priority to an ephemeral food plate (representing a client) over a permanent plate doubled the food reward (Bshary and Grutter, 2002; Triki *et al*., 2019). To understand if cleaner gobies are able to solve partner choice cognitive tests, we tested captive-bred dedicated cleaner gobies *Elacatinus oceanops*, commonly known as neon goby, for their capacity to solve a biological market test. We varied cue-based decision-making using both ecologically relevant cues (plate size and colour) and non-relevant ones (presentation side). Notably, size and colour are considered ecologically relevant cues because they mirror essential client characteristics, such as species identity. In their natural habitat, cleaner gobies rely on visual cues like size to identify clients with higher ectoparasite loads, which is crucial for their feeding, providing efficient cleaning services and shaping decision-making strategies. Additionally, we tested their capacity for reversal learning, an indicator of complex cognitive abilities.

## 2 Material and methods

### 2.1 Study species and experimental design

Cleaner gobies, *Elacatinus oceanops*, were bred in an ornamental aquaculture facility and transported to the aquatic facilities of Laboratório Marítimo da Guia (Cascais, Portugal) by TMC-Iberia. A total of 13 cleaner gobies were individually housed and acclimated to the experimental setup by feeding them mysid shrimps spread on Plexiglas plates.

The study took place from November 2022 to January 2023. We adapted our experimental design from the biological market theory experiment introduced by Bshary and Grutter in 2002. This design has been widely replicated (Wismer *et al*., 2014; Paula *et al*., 2019; Triki *et al*., 2019) and further modified by Wismer et al. (2019). In our setup, individual aquariums were divided into two sections by a transparent partition (see Fig. 1). The fish were kept in the smaller section, while experimental Plexiglass plates were introduced into the larger section. When trials began, the partition was removed, giving the gobies access to two Plexiglass plates, each with a small amount of mysid shrimp (approximately 0.001 g). One plate was ephemeral and removed if not chosen first, while the other was permanent, remaining in the aquarium for the entire duration of the trial (around 2 min.), regardless of the gobies’ foraging decisions.

**Figure 1.**
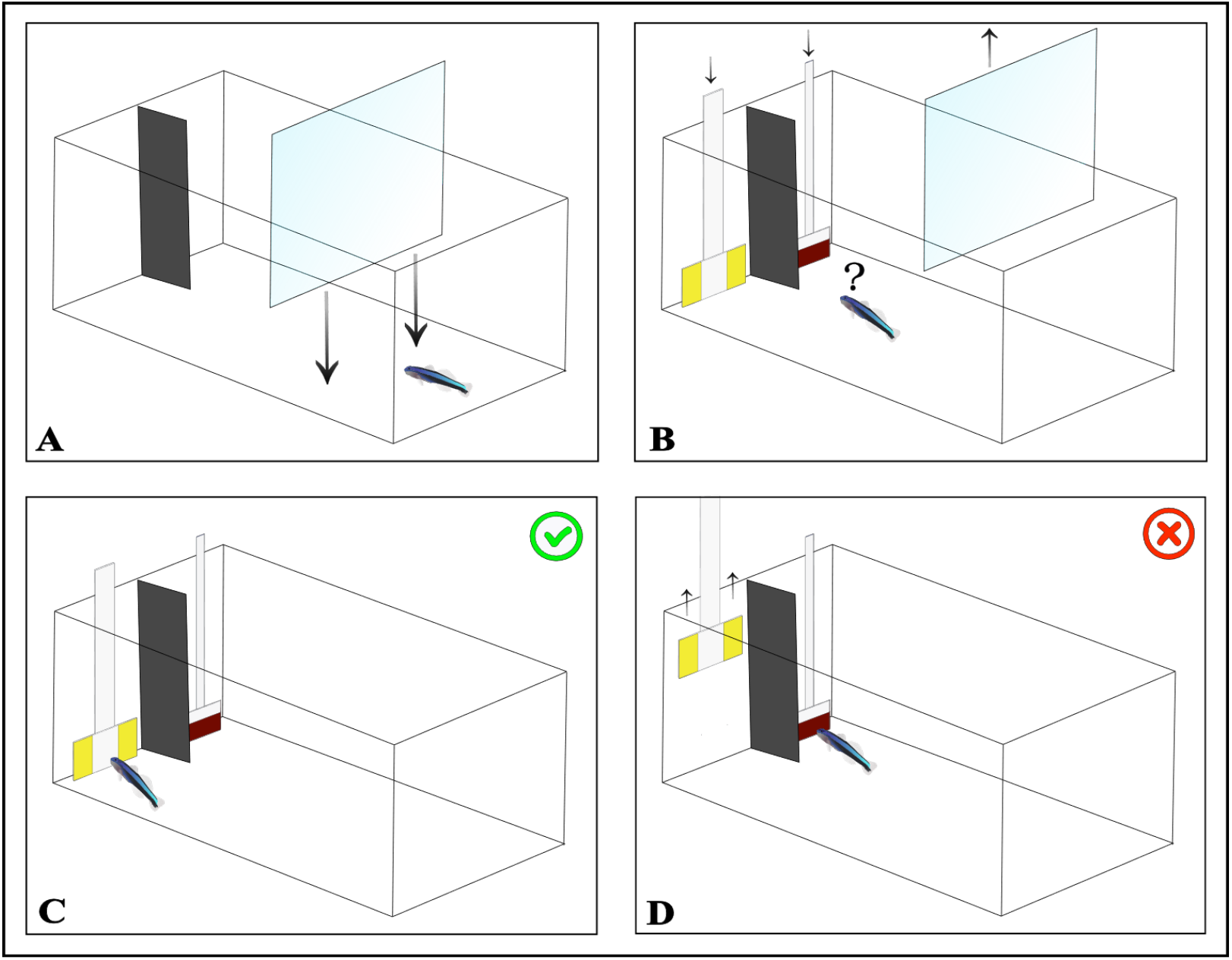
Test procedure. A) The fish was initially guided into one side of the aquarium using a transparent partition; B) Subsequently, the transparent partition was removed, allowing the fish to forage freely; C) In the event of the fish choosing the correct plate, both plates remained in the aquarium, granting access to the food reward; D) Conversely, if the fish opted for the incorrect plate, the opposing plate was promptly removed, denying access to the reward.

In this study, the established protocol was followed with one key difference: the method of indicating which plate was ephemeral varied by test, using either size, colour, or spatial cues. For all tests except those involving spatial cues, the placement of the ephemeral plate in the aquarium (either left or right) was balanced and semi-randomized across 10 trials. A lottery system ensured the ephemeral plate appeared on each side 50% of the time, without being on the same side more than three times consecutively. The number of trials it took for the cleaner fish to learn to feed from the ephemeral plate first was monitored. The learning criterion was deemed met if the cleaners chose the ephemeral plate first in at least 9 out of 10 consecutive trials, 8 out of 10 trials in two consecutive sessions, or 7 out of 10 trials in three consecutive sessions. Each cleaner fish was exposed to all cue types in the same order: plate size, then a reversal, followed by colour, another reversal, spatial cues, and a final reversal. Each fish participated in one session per day, with each session comprising 10 trials, totalling 100 trials per cue type.

#### 2.1.1 Test 1: Variation in size

Two Plexiglas plates, one large (7 cm x 4 cm) and one small (4 cm x 3 cm), characterised by two different colours and patterns (i.e., two yellow vertical stripes and one brown horizontal stripe), were used. The larger plate represented the ephemeral plate (correct choice), and the small one represented the permanent (wrong choice). The presentation side of each plate size was balanced and randomised across the 100 trials.

#### 2.1.2 Test 2: Variation in colour

The second test started 60 days after the first. During the 60 days, fish were fed with white plexiglass plates without any colour or size information to desensitise any learning patterns from the first test. Plates of equal dimensions were presented in colours and patterns matching those of Test 1. The plate positions (left or right) were balanced and randomised across the 100 trials.

#### 2.1.3 Test 3: Spatial cue

The test involved associating a specific location (left or right side of the tank) with the food reward. Two white Plexiglas plates of equal size were used. The correct plate location (left or right) was randomized, as in Tests 1 and 2.

#### 2.1.4 Reversal Learning

Following a 12-days break after each cue test, the fish that successfully completed the initial test were subjected to a reversal test. In these tests, the cue associated with the ephemeral plate was switched for each type of cue: size, colour, and spatial. The procedures remained the same, except for a key change: the behaviour of the plates was reversed. The plate that was previously ephemeral (removed if not chosen first) now remained in the tank permanently, regardless of the fish’s choice, while the plate that was previously permanent became the ephemeral one. This reversal was implemented abruptly to assess the fish’s adaptability to changes in cue associations.

The colours displayed in the figure are the same as those used in the experimental procedure.

### 2.2 Time frame of the tests

Test 1 was performed between 17 November and 26 November. Reversal of Test 1 was performed 25 days after Test 1, between 13 December and 22 December 2022. Test 2 was performed between 17 January and 26 January 2023. Reversal of Test 2 was performed 12 days after Test 2, between 30 January and 8 February 2023. Test 3 was performed between 14 February and 23 February 2023. Finally, reversal of Test 3 was performed 12 days after Test 3, between 27 February and 8 March 2023.

### 2.3 Statistical Analysis

Survival analyses were conducted for each test using the R package “survival” (Therneau, 2015) to analyse time event data (number of trials to solve the task) and compare tests with different ecological relevance for the study species. The survival analysis was fitted using the Cox proportional hazard model (‘coxph’). Initially, a survival standard object was constructed (‘surv’), and subsequently, to assess statistically significant differences in the learning performances among the tests, a non-parametric long-rank test was performed (‘survdiff’). This test allowed us to compare the survival curves representing the task completion over time. To validate the assumption of proportional hazards, a Schoenfeld test (‘cox.zph’) was conducted. Following this, the residuals were graphically examined (‘ggcoxzph’) to identify any time-dependent effects of the ‘correct’ variable on the hazard.

## 3 Ethical Note

Animal experimentation met the ASAB guidelines for the ethical treatment of animals. The experiments were conducted under the approval of Faculdade de Ciências da Universidade de Lisboa animal welfare body (ORBEA-FCUL, ref: 04/2023)

## 4 Results

Out of the initial 13 *E. oceanops* individuals, 13 were tested for size cues and 12 for colour and spatial cues, as one fish died during the experiment. Among them, 10 (77%) successfully solved the task using the plate size as a cue (Fig. 2A, χ^2^=11.55, p=0.003). Subsequently, when presented with both colour/pattern and spatial cues, only 2 (17%) passed the test. Notably, none of the gobies successfully solved the reversal version of any cue (Fig. 2B).

**Figure 2.**
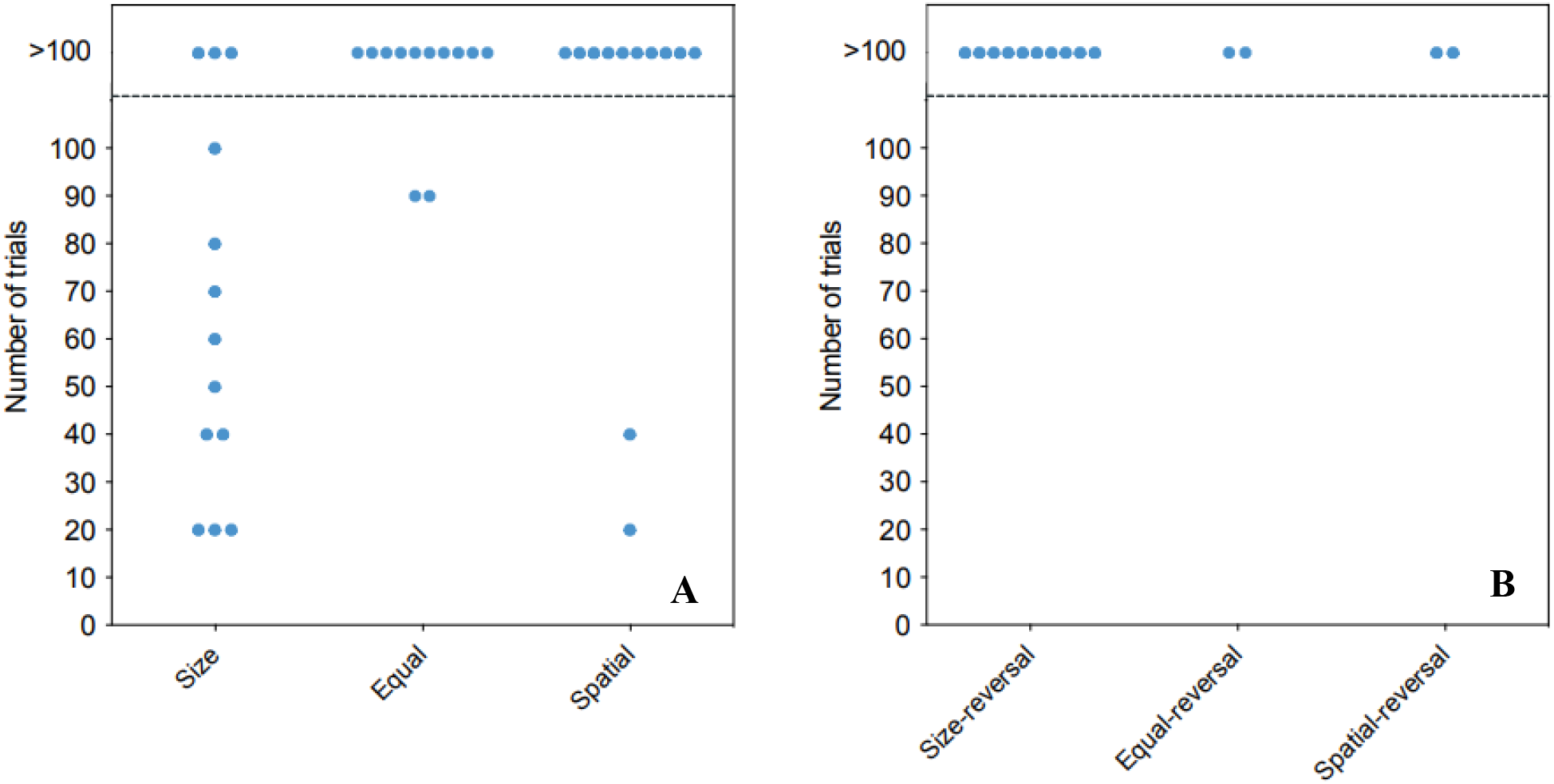
Performance of cleaners in the cognitive tests. (A) The number of trials required for cleaners to establish a significant preference for the visitor plate. Each circle represents the performance of one individual. Individuals above the dashed line did not complete the task within the maximum of 100 trials. Cue type, namely size, colour and spatial, is represented on the y-axis. (B) The number of trials required for successful cleaners to reject their established preference and learn to prefer the reversal cue.

The survival analyses, conducted using the Cox proportional hazard model, revealed significant differences in learning performances among the tests (χ^2^=15.8, p<0.0001). Moreover, the analyses revealed significant time-dependent effects of the “correct” variable on the hazard, suggesting deviations from the proportional hazard assumption (Schoenfold test, p<0.05).

When individuals were tested with the same size plate, and during the spatial discrimination test, they exhibited significantly lower hazard rates compared to those tested with plates of different sizes (p=0.00734; p=0.01114, respectively).

**Table 1.** Sample size and experimental conditions. Number of fish tested for three different tasks and reversal learning: “Variation in size” (Test 1), “Variation in colour” (Test 2), and “Spatial cue” (Test 3).

| Number of fish tested |  |  |  |  |  |  |
| --- | --- | --- | --- | --- | --- | --- |
|  | Test 1 Variation in size |  | Test 2 Variation in colour |  | Test 3 Spatial cue |  |
| Species | BM | BM Reversal | BM | BM Reversal | Spatial cue | Reversal |
| *E.oceanops* | 13 | 10 | 12 | 2 | 12 | 2 |

## 5 Discussion

Understanding the mechanisms behind the diversity in decision-making rules is crucial for grasping how natural selection shapes social behaviours in response to environmental needs. Our study focused on whether cleaner gobies, *Elacatinus oceanops*, could successfully complete three discriminatory two-choice tasks in a biological market scenario by employing various cue-based decision strategies. We observed that cleaner gobies could successfully accomplish this test when specific cues, particularly size, were utilised but failed when cues such as colour were included. These findings emphasise the critical significance of accurately identifying salient cues to test cognitive performances.

Previous studies on cleaner wrasse (*Labroides dimidiatus*) and capuchin monkeys in biological market task variations highlighted the importance of cue type for solving these tasks (Salwiczek *et al*., 2012; Prétôt, Bshary and Brosnan, 2016; Triki *et al*., 2019a). Similarly, our findings confirm the significance of cue type in determining the ability of cleaner gobies to complete a biological market task. When size was the cue, a significant majority of cleaner gobies (77%) successfully solved the test by choosing the larger plate, indicating the universal recognition of size as an informative cue. In their natural habitat, size is a practical cue, as larger fish typically host more ectoparasites (Grutter, 1995; Grutter and Poulin, 1998), and gobies show a preference for clients with a higher parasite load (Sikkel, Cheney and Côté, 2004; Soares, Cardoso and Côté, 2007). Additionally, observations revealed that gobies prioritise predator clients, often characterised by larger body sizes. Prioritising predators aids gobies in reducing the negative impact of a potential threat to the other fish, encouraging the clients to return promptly, and facilitating the mutualistic exchange of benefits (Soares, Cardoso and Côté, 2007). Another important aspect is the cleaning dependence of the species study. *Elacatinus oceanops* is a dedicated cleaner, exhibiting behaviour that requires sophisticated mechanisms. Their ability to solve the test where the decision-making criteria are associated with the size might reflect the precision with which their cognitive skills have adapted to meet their needs. This not only highlights the adaptability of their cognitive abilities but also underscores the significant role played by the feeding behaviour for which they were naturally selected.

However, in our study cleaner gobies struggled to use colour cues effectively. This unexpected result contradicts previous assumptions about the importance of colour in species recognition during cleaning interactions. While parasite loads and mucus quality vary not only with the size of the fish but also across different species (Grutter, 1994; Arnal, Côté and Morand, 2001), our findings suggest a reliance on size rather than colour or patterns in the decision-making process of cleaner gobies. This result underline the importance of the size as a salient cue in the learning process of this fish, also considering the observation in an experiment conducted with a closely related cleaner goby species, *Elacatinus prochilos* (Mazzei *et al*., 2019), where gobies exhibited limited learning abilities to discriminatory tasks involving different patterns but same colours and size. The reliance on size cues over colour or pattern cues may reflect the importance of the physical characteristics of clients.

Furthermore, in contrast to their proficiency with ecologically relevant cues like size and their related species *E. prochilos*, cleaner gobies displayed a notable lack of cognitive ability in a test unrelated to their natural ecological context, the spatial cue (Test 3). In this test, the fish were required to distinguish between spatial arrangements to successfully complete the test. Unlike the tests involving size cues, where the gobies could rely on their innate or learned ecological behaviours, the spatial test demanded a different kind of cognitive processing. We propose two possible explanations for the observed lack of flexible cognition. Firstly, the spatial discrimination test likely posed a challenge because it required the gobies to utilise spatial memory skills, which are not as directly linked to their natural behaviours. In their natural habitat, cleaner gobies are more likely to rely on visual cues like size and colour to identify clients with higher ectoparasite loads, which is crucial for their feeding and survival. However, navigating and remembering spatial configurations does not have a direct bearing on these essential tasks. Another plausible explanation could be a consequence of the experimental procedure. The fish were exposed to different tasks over a period of three months. This prolonged training period might have resulted in decreased motivation and a potential failure in successfully navigating the spatial discrimination task.

While these findings may indicate low cognitive flexibility in cleaner gobies, it is essential to note potential confounding factors, including using fish raised in captivity. Studies have shown that individuals raised in not complex environments may exhibit different and lower cognitive performance than their wild counterparts raised in complex and enriched environments due to a lack of environmental experiences (Wismer *et al*., 2019). Furthermore, the importance of testing species in their natural habitat is underscored by previous research (Bräuer *et al*., 2020; Salena *et al*., 2021), highlighting the impact of artificial selection and life experiences on cognition.

Given the potential variance in the impact of natural selection between laboratory and wild environments, the observed low capacity in cognitive abilities and the absence of flexibility may be attributed to biases inherent in laboratory conditions. Moreover, the experimental setup, notably the two-week gap between tasks, and potential sequence effects might have impacted the strength of the association between stimuli and reinforcement. This choice might have influenced the performances of the fish. Therefore, the apparent challenges in task-solving proficiency could arise from factors, such as prolonged training regimens, diminished motivation, and stress induced by the test procedures. Furthermore, the inability to learn subsequent discriminations does not necessarily signify inferior cognitive capacity, as environmental and motivational variables can impact task execution.

However, the fact that these gobies demonstrated cognitive skills in tests closely resembling their natural ecological scenarios indicates an inherent, specialised learning mechanism. This tendency of cleaner gobies to effectively process and respond to cues relevant to their natural behaviours, despite their lack of direct environmental experience, emphasises the potential for a latent cognitive framework shaped by evolutionary history. While it’s understood that animal cognition is shaped by a complex interplay of various factors, including both biotic and abiotic elements, genetics, and evolutionary processes, our findings particularly highlight the significant role of evolutionary influences. This underscores the profound impact of evolutionary history on shaping innate cognitive abilities in animals, particularly in relation to ecological relevancy.

While our findings hold significance on the innate cognitive skills of cleaner gobies, further research incorporating wild populations is essential to comprehensively understand their holistic cognitive abilities. Studying these fish in a laboratory setting, especially those reared individually, primarily sheds light on their innate cognitive skills. However, extending this research to include wild populations of cleaner gobies could offer additional insights and a deeper understanding of the species’ cognitive capabilities. Conducting similar tests in the wild would not only help determine if their ability to utilise other cues, such as colour or patterns, could be enhanced through natural cleaning interactions, but it would also provide a more holistic view of their cognitive skills. In their natural habitats, cleaner gobies must continuously adapt their behaviour to the ever-changing environmental conditions.

In conclusion, our study highlights the critical role of incorporating cue salience within an ecological context for understanding the variations in social decision-making processes. By examining how specific cues influence cognition, we can better appreciate the intricate balance between innate abilities and learned behaviours in shaping social decision rules. This approach not only enriches our comprehension of the evolutionary and environmental factors driving these variations but opens up new avenues for exploring the complex interplay between an organism’s innate predispositions and its adaptive responses to ecological demands.

## 8 Acknowledgments

We would like to acknowledge all members of the Behavioural Ecology and Evolution group who contributed with comments and help throughout this work.

## Data Availability Statement

The datasets generated for this study can be found in FigShare [https://figshare.com/s/03356c39dad82e9dc6ba].

